# *Sporosarcina pasteurii* can form nanoscale crystals on cell surface

**DOI:** 10.1101/184184

**Authors:** Swayamdipta Bhaduri, Tanushree Ghosh, Carlo Montemagno, Aloke Kumar

**Author notes:** Equal contributions (Swayamdipta Bhaduri, Tanushree Ghosh).

## Abstract

Using a semi-solid 0.5% agar column, we study the phenomenon of microbially induced mineral (calcium carbonate) by the bacteria *Sporosarcina pasteurii*. Our platform allows for *in-situ* visualization of the phenomena, and we found clear evidence of bacterial cell surface facilitating formation of nanoscale crystals. Moreover, in the bulk agar we found the presence of microspheres, which seem to arise from an aggregation of nanoscale crystals with needle like morphology. Extensive chemical characterization confirmed that the crystals to be calcium carbonate, and two different polymorphs (calcite and vaterite) were identified.

## Introduction

Biomineralization refers to the process of mineral precipitation due to chemical alteration of the environment induced by the microbial activity^1–4^. For unicellular organisms such as bacteria^4^, the biomineralization process can be either extracellular^5^ or intracellular^6^. Microbially induced calcite precipitation (MICP) is an excellent example of an extracellular mineral deposition. Several microbial species take part in MICP by means of various mechanisms such as photosynthesis^7,8^, urea hydrolysis^1,5^, sulfate reduction^9,10^, anaerobic sulfide oxidation^11^, biofilm and extracellular polymeric substances^12,13^. There has been significant interest in microorganisms that can produce urease (urea amidohydrolase; EC 3.5.1.5) and hence are able to hydrolyze urea to induce calcite precipitation^14^. *Sporosarcina pasteurii* (formerly *Bacillus pasteurii)* is a non-pathogenic, endospore producing soil bacteria that also produces urease and can tolerate highly basic environments. *S. pasteurii* has attracted significant attention from researchers for its unique feature of calcium carbonate precipitation, which can be easily controlled^10,15–19^. *S. pasteurii* is being investigated for the possibility of its utilization towards a multitude of applications including underground storage of carbon, healing masonry structures of archaeological importance and long-term sealing of geologic cracks in large-scale structures^3,5,20–24^ Hammes and Verstraete (2002) reported the four very important parameters for MICP as pH, dissolved organic carbon, calcium concentration and available nucleation site^14^. The saturation rate of carbonate ions concentration (CO_3_^2-^) is controlled by the first three parameters and it is believed that bacterial cell wall as the nucleation site facilitate the stable and continuous calcium carbonate deposition^20^. However the bacteria-free solution loaded with urease enzyme can also induce calcium carbonate precipitation. Mitchell and Ferris (2006) studied the influence of bacteria on the nucleation of MICP. The bacteria-free enzyme solution was compared to the bacteria induced environment. Authors reported significant positive effect of bacterial presence (referred as “bacteria-inclusive”) on the increase of size and growth rate of the precipitated crystal though the idea of bacterial control of MICP was rejected. The bacteria-free urease solution showed similarities with the bulk chemical precipitation whereas differences were observed for aqueous microenvironment of bacteria^16^.

It has been suggested that the bacterial cell walls can serve as nucleation sites^3,14,23,25^ as bacterial cell surfaces carry negatively charged groups at neutral pH^26–28^. These negative charges can influence the binding of cations (e.g. Ca^2+^) on the cell surface and eventually act as nucleation sites when reacting with carbonate anions from the solution to form insoluble calcium carbonate^1,26,29^. Definitive proofs of involvement of bacterial cell walls in complex environments on the nucleation process are rare. Ghashghaei and Emtiazi reported the presence of nanocrystals of calcium carbonate on cell walls of the bacteria *E. ludwigii* for experiments performed with a liquid culture^30^. While the data is suggestive of the role of cell wall, a more definite proof is desirable. In a contrasting study, Bundeleva et al. reported on MICP by the anoxygenic *Rhodovulum* sp. Their TEM analyses failed to show the existence of CaCO_3_ on or near live cells^31^. The authors claim the existence of certain cell protection mechanisms against mineral incrustation at the vicinity of live bacteria and they invoke the idea of mineral precipitation at a certain distance from the cell surface^31^. Thus, we see that the issue of role of cell wall on mineral precipitation in MICP remains controversial and detailed studies delineating the exact mechanism leading to the nucleation of crystals in *in-situ* conditions are required to resolve this important biophysical conundrum.

To the best of our knowledge, no conclusive reports of the role of cell wall in MICP in complex environments have been reported. Applications of *S. pasteurii’s* MICP often include their activity in porous media such as concrete and rocks^3,20,32–34^, where *in-situ* visualization is often challenging. In order to the address the fundamental mechanism of MICP by *S. pasteurii* in complex environments, we designed an agar column setup, which allows us to investigate mineral precipitation by direct observation. Our setup consists of a 0.5% agar-column that is stab-inoculated and MICP proceeds in the setup as downward travelling conspicuous mineral trails. We investigate the morphology of precipitated crystals and conclusively demonstrate that the presence of nano-crystals of calcite on the bacterial cell wall. While this can be considered as definitive evidence that the cell wall does serve as a nucleation site, other mechanisms of nucleation are not ruled out.

## Results and Discussion

### Agar Column

To understand growth of *S. pasteurii* is a porous environment, an agar column with stab culture was observed for a period of one week. Figure 1(a) shows the original culture tubes containg growth spans after day-1 and day-7. Extensive calcite deposition can be prominently seen in Figure 1(b) after 7 days of incubation. The agar media acted as soft, porous and transperant nutrient riched environment to monitor the bacterial motility and calcite precipitation. Figure 1c depicts a scanning electron microscopy image of the agar column, where porous structures can clearly be seen. *S. pasteurii* bacterium possesses flagellum (see Figure S1b) which likely allows it to navigate the porous structure of the agar column. The agar column was sectioned at a depth of 1.25 cm after 4 days, and the bacterial motion observed through optical microscopy. Figure S2 a,b show a motile bacteria within the agar column and the corresponding velocity historgram. The histogram indicates within the agar matrix the bacterium can travel at velocities ~10^−7^ m/s and hence it can be expected that bacterial cells can travel approximately 1 cm of agar column per day. This ‘seeping speed’ is also corroborated through macroscale observations (Figure 1b). Figure 1d also shows the presence of deposits in the agar column after 7 days of bacterial activity.

**Figure 1.**
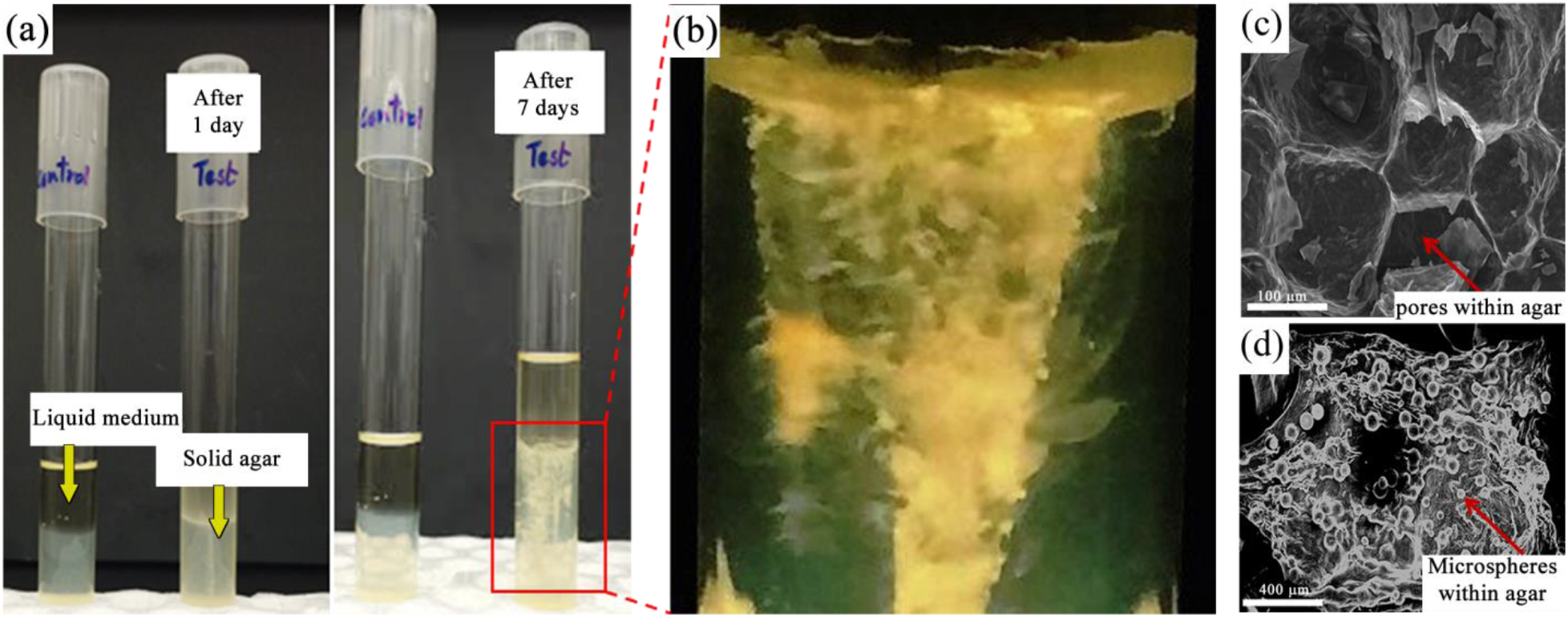
(a) Two samples containing the semi-solid 0.5% agar medium in two identical 20 mL test tubes. The one to the left is the control sample which is devoid of any bacterial cells. To the right is the *S. pasteurii* inoculated sample where the resulting mineral precipitation has left a conspicuous trail. Images were taken after 1 day and 7 days of the inoculation. (b) Blown out magnified image of the mineral deposition. (c-d) SEM image of agar showing pores in control set (c) and numerous mineral microspheres after 7 days (d) of incubation.

### Calcite micro-spheres

The primary advantage of the agar-column experimental setup is *in-situ* visualization and easy access for supporting characterizations. The agar column was section at a depth of 2.5 cm after a period of 7 days and crystalline depositions were characterized by other forms of microscopy. Specifically, the agar colunm was characterized using optical microsopy, transmission electron microscopy (TEM), selected area diffraction pattern (SAED) and elemental characteization was determined by powder XRD and energy-dispersive X-ray spectroscopy (EDS). Optical microscopy of ultrathin (~80 nm) sections of the agar medium revealed depositions of crystalline mircrospheres profusely within the agar column. Optical and TEM imaging revealed that the visible white deposits within the column were mostly crystalline micropheres of about 10-50 µm in diameter. Figure 2(a) shows a crystal violet stained corss-section of microsphere (diameter ~10 µm) and Figure 2b shows the TEM image of the same with a SAED pattern (Inset of Figure 2(b)) proves the crystalline nature of the precipitates and also consistent with the Miller indices of calcium carbonate observed in XRD. The detailed magnified observation of the microspheres in Figure 2 (c and d) revealed the spatial arrangement of calcite nano-needles at the interface of agar-calcite microsphere. Elemental confirmation was performed with EDS on selected microsperes at the agar-crystal interface (Figure 2c) and within the microcrystal (Figure 2d). EDS (Figure 2e) spectral results indicated the presence of Ca, C and O. The direct *in-situ* visualization on the MICP provides a new window to explore the arrangement of calcium carbonate nano-needles, which give rise microspheres within the semi-solid porous agar media.

**Figure 2.**
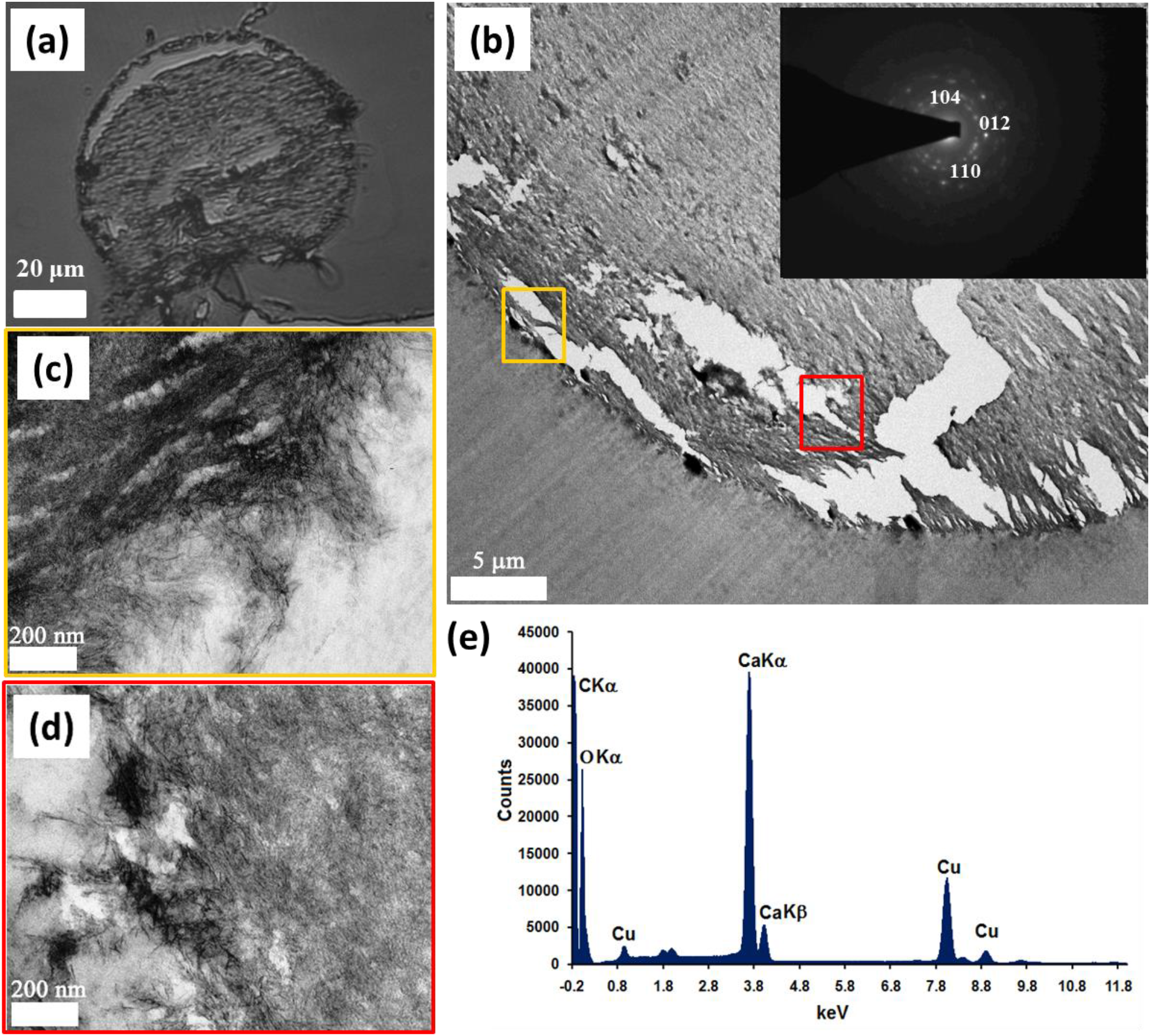
(a) An optical microscopy image of an ultrathin (80 nm) section of agar column with crystal, which was stained with crystal violet. (b) TEM image of a portion of the same crystal. (inset) The SAED pattern of the crystalline structure. (c)-(d) Higher magnification TEM images of the crystal shown in (b). Corresponding locations are boxed and color coded (e) The EDX analysis indicates the presence of C, Ca and O.

To further confirm the elemental distribution, the dissected calcium carbonate microspheres (diameter ~50 μm) were identified within the freeze-dried agar column and subjected to secondary electron SEM (Figure 3a) and backscattered SEM (BSE) (Figure 3b). The increased intensity in BSE image (Figure 3b) confirmed the presence of highly dense elements with higher atomic number e.g. Ca. EDS elemental mapping (with false colors to different elements) also supported the compositional presence of Ca (Figure 3c) along with other pertinent elemental compositions of the microspheres to be C (Figure 3d) and O (Figure 3e) were once again verified. The Cl mapping (Figure 3f) showed negligible signal on the surface whereas prominent signal from the dissection. This signal is likely due to the added NaCl in the bacterial growth media. The transparent agar column therefore provided a real time monitoring platform to study *S. pasteurii* induced calcium carbonate precipitation. The optical and elemental confirmation of the precipitate provides an experimental support to the mineral deposition phenomenon induced by bacteria.

**Figure 3.**
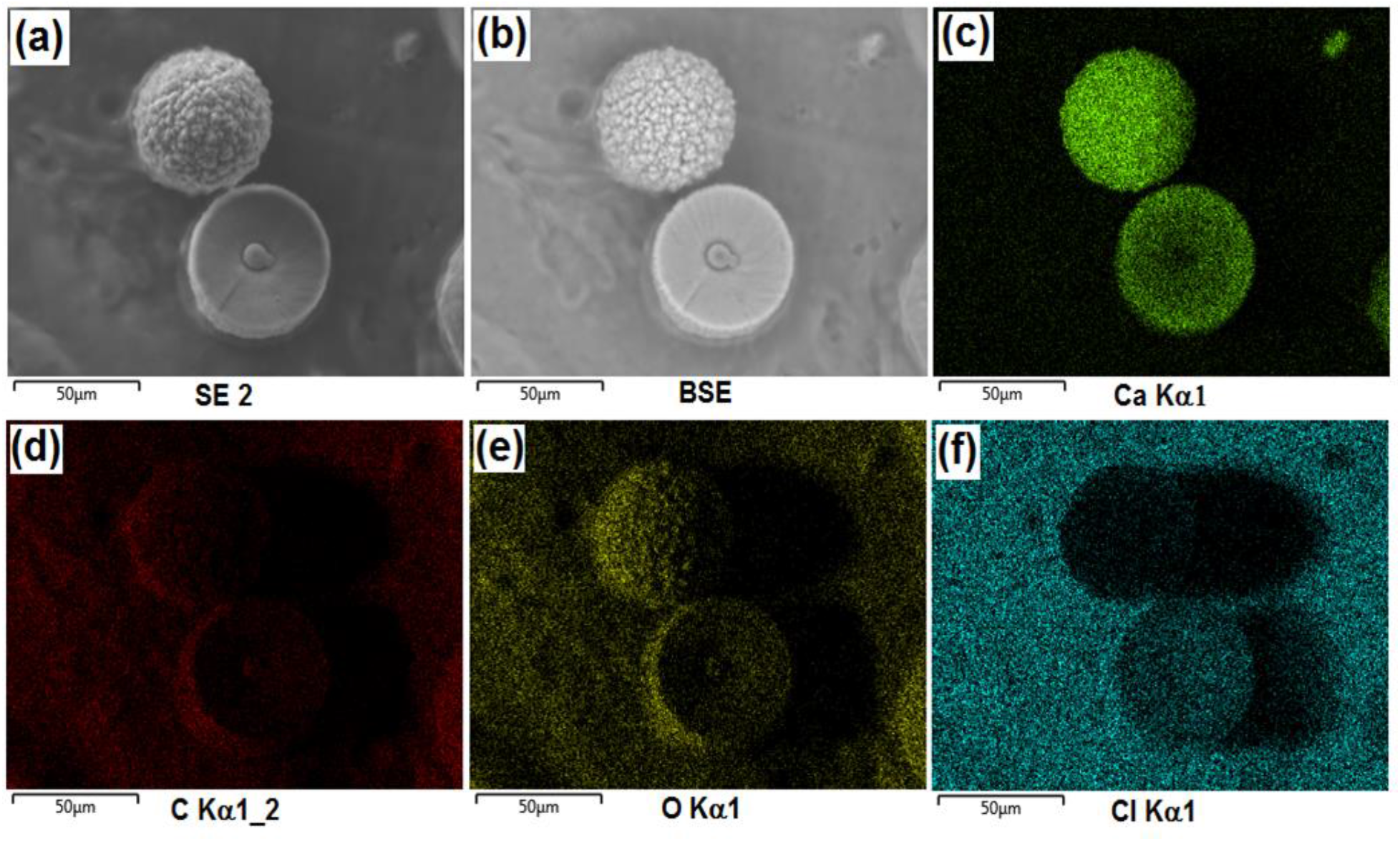
SEM and EDS mapping of elemental distribution analysis for calcite microspheres. (a) Secondary electron microscopy (SE 2) (b) backscattered electron microscopy (BSE) images. (c-f) EDS mapping of elemental distribution showing Ca (c), C (d) and O(e) as the main ingredient of microspheres. Appearance of Cl (f) within microspheres and surrounding agar possibly originates from added NaCl salt in the bacterial growth medium.

### *S. pasteurii* micro-environment

TEM imaging of the micro-environment of bacterial cell surface shed light on the likely nucleation route for the calcite microspheres. The 80 nm ultra-thin sections of agar column containing embedded bacteria shows in Figure 4(a, b and c) clearly depict the micro-environment of the bacterium. Oval shaped bacterial cells can be seen along with formation of spores (Figure S1a). Nano-needles that were previously encountered (Figure 2c) also seen at the vicinity of the cells (Figure 4a). Magnified images indicate nanoscale spherical depositions on cell surface (Figure 4b) and needle-like depositions in the surrounding agar media (Figure 4c). The elemental confirmation was performed with EDS on bacterial cell surface deposition. EDS results (Figure 4d) indicated the presence of Ca, C and O. The powder XRD results of the whole colunm used to identify the crystalline phases of the inorganic compounds. The identified signature peaks of calcite (Figure 4e) at 2θ valus of 26.83°, 34.26° and 42.01° respectively corelated with lattice (hkl) indiced of (012), (104) and (110). Low intensity vaterite peaks (Figure 4e Inset) were also identified at 2θ valus of 28.98°, 31.52° and 38.22° corelated with lattice (hkl) indiced of (100), (101) and (102), respectively. With these results, we can definitely conclude that the highly elelctron dense depositions on the bacterial cell wall were actually calcium carbonate crystals. XRD and EDS results strongly support the existance of calcium carbonate and its polymorphs as nanoscale deposits on the bacterial cell wall. These results are definitive proof that the bacterial cell wall does participate in the MICP process as a nucleation site.

**Figure 4.**
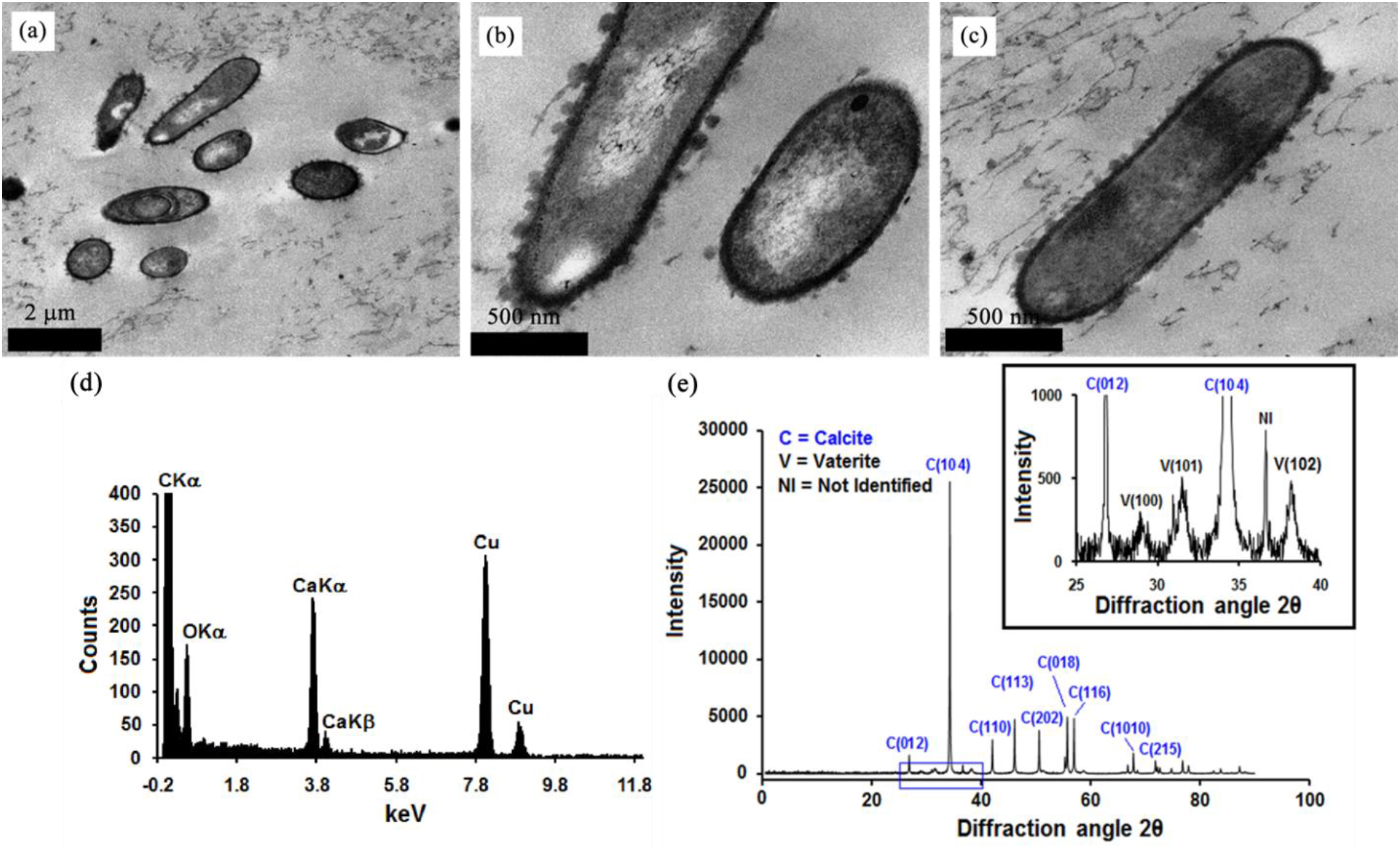
(a)-(c) TEM images of ultrathin sections of the agar medium depicting the presence of cells and spores. They also show the presence of bacterial cell surface depositions. (d) EDS pattern of cell surface deposition shows existence of Ca, C and O. (e) XRD plot of the same location indicates the presence of calcite and vaterite polymorphs. (inset) Blown-up sub-section of (e) clearly demonstrates presence of vaterite.

## Discussion

In this work, we develop a simple, yet effective experimental platform for *in-situ* imaging of the MICP process by *Sporosarcina pasteurii*. Agar (0.5%) was used to create a porous column, which was inncoluated with a stab culture. As *S. pasteurii* cells migrated down the agar column they left conspicuous trail of crystals. Samples of the agar column at different locations were taken and subjected to microscopy and we found that the crystal train consisted of calcite microspheres, which on closer inspection were found to be an aggregate of needle-like nanoscale crystals. Moreover, cells whose surface contained calcite nanocrystals were also observed confirming the hypothesis that cell surface plays a role in nucleation.

## Materials and Methods

### Bacteria culture

The bacterial strain *Sporosarcina pasteurii* (Miquel) Yoon et al. ATCC^®^ 11859^™^ was obtained from American Type Culture Collection (ATCC) in freeze-dried condition. The strain was first cultured in ATCC recommended medium^1^. S. pasteurii cultures were further prepared in nutrient media^35^. Nutrient Broth (NB) was prepared, which contained 5 g peptone; 3 g beef extract and 2 g sodium chloride per 1L of distilled water. The pH of this medium was adjusted to 7.0 using HCl and NaOH. Nutrient agar medium was prepared by utilizing the same ingredients as NB with an additional supplement of 1.5% bacteriological agar. Solution mixtures were sterilized by autoclave wet sterilization method (121 °C, 15 psi) for 15 min. NB and NB-agar were subsequently used for suspension culture and sub-culturing of S. pasteurii before the experiments. All the cultures were incubated in aerobic conditions at 30 °C. Peptone, beef extract, urea, NaCl, HCl, NaOH and bacteriological agar were purchased from Fischer scientific (Thermo-Fisher Scientific, Waltham, Massachusetts, USA). CaCl_2_. 2H_2_O (ACS reagent ≥ 99%) was purchased from Sigma-Aldrich (Sigma-Aldrich, St. Louis, Missouri, USA). All the chemicals were used as purchased and solutions were prepared in Milli Q (18.2 MΩ) water.

Preparation of semisolid-agar column: Semisolid-agar columns were prepared by following Bang’s urea-CaCl_2_ liquid media with modification^1^. The modified media contain peptone 0.5 %; beef extract 0.3 %, sodium chloride 0.2 % and CaCl_2_, 0.28 %. The pH of the medium was adjusted to 7.0 and then 0.5 % agar was added prior to autoclaving. Urea (2 %) was added separately after autoclaving when media temperature cooled down to approximately 50-60° C. To create the agar columns, 10 ml of liquid agar was poured into upright test tubes and allowed to cool inside a biosafety cabinet, which finally resulted in columns of approximately 5 cm in length. Subsequently, these agar columns were inoculated by stabbing the free surface of the agar column with pre-cultured S. pasteurii using a stabbing needle. Fresh liquid media were poured on to the agar column to prevent drying out from the agar surface. The bacteria inoculated columns along with a control were incubated at 30 °C for a maximum duration of 7 days.

### Microscopy

For microscopy, sample volumes were cut from the agar-column approximately at mid-height (~ 2.5 cm). Agar slices were cut into small cubes and fixed in solution containing 2.5% glutaraldehyde, 2% paraformaldehyde in 0.1 M phosphate buffer (pH=7.4) for 30 min. The fixation process was followed by buffer wash (0.1M phosphate buffer). Post-fixation treatment of 1% Osmium tetroxide in 0.1 phosphate buffer was performed for 1 h. Standard protocols for buffer washing and dehydration through graded alcohol were followed^36^. Samples were infiltrated with spurr resin (1:1 of Ethanol: Spurr mixture) for 3h and then kept in 100% spur for 24h embedding. Samples were embedded in flat molds with fresh spur and cured at 70oC for overnight. Cured embedded resin capsules were sectioned in 70-90 nm thin sections using ultramicrotome (Reichert-Jung UltraCut E, Vienna, Austria) and mounted on copper grid for transmission electron microscopy (TEM, Philip-FEI, Morgangni 268, Oregon, USA) operated at 80kV.

Bacterial cultures grown in urea-CaCl2 liquid and solid media were observed for calcium carbonate deposition using scanning electron microscopy (SEM, Zeiss EVO M10, Oberkochen, Germany) and optical microscopy (Nikon Eclipse Ti, Nikon Instruments Inc., Melville, USA). The SEM samples were prepared by fixing and dehydration in graded alcohol similar to TEM. Dehydrated sample were kept in 100% ethanol and mounted up on carbon tape. The crosssections of the agar-column were freeze dried (SuperModulyo Freeze Dryer, Savant Instruments Inc., New York, USA) after fixation and directly used for SEM imaging. Every sample was gold sputtered (Denton Vacuum, Desk II, MOORESTOWN, New Jersey) before SEM imaging.

For cell motility experiments, two parallel setups were used – liquid and semi-solid agar media. The liquid media experiment was set up in a Lab Tek II Chambered Coverglass Slide (Thermo Fisher Scientific, Waltham, MA) and agar media experiment was conducted by taking a slice from the agar column at approximately mid-height. Images were obtained using the Nikon Ti inverted microscope using the bright field mode. The camera used was Nikon DS-Qi1Mc, which was operated at 10 fps. Image analysis was performed using ImageJ.

## Acknowledgements

This study has drawn heavily on instrumentation like SEM, TEM, SAED, EDX and XRD. All microscopy and characterization technicians at the NanoFab and Department of Biological Sciences, University of Alberta are thankfully acknowledged.

## Author contributions

TG and SB performed experiments, analyzed data, prepared results and wrote the manuscript. TG and SB shared equal contributions. AK conceived part of the work and analyzed data. AK and CM supervised the overall research and participated in manuscript writing.

